# Microbiota-driven, strain-specific *Asaia* colonisation modulates mosquito vector competence

**DOI:** 10.64898/2026.09.09.750370

**Authors:** Bahrami Romina, Di Castri Sofia, D Gingell Daniel, Cappelli Alessia, Favia Guido, Damiani Claudia, Mancini Maria Vittoria

## Abstract

Microbiota-based mosquito control strategies are founded on the ability of bacterial symbionts to modify host biology and/or pathogen transmission. However, symbionts do not operate in isolation. Their colonisation, persistence and biological effects arise within a complex ecological network comprising the host and its resident microbiota. We hypothesized that colonisation success is not only an intrinsic property of a bacterial symbiont but emerges from interactions between bacterial strain, host background, and the native microbial community. To test this, we compared recolonisation by endogenous and exogenous *Asaia* strains in the invasive mosquito *Aedes koreicus* using wild-type and axenic mosquitoes to experimentally manipulate the microbial composition. We found that endogenous and exogenous strains followed distinct colonisation trajectories. Depletion of the resident microbiota selectively rescued colonisation by the endogenous strain, whereas both strains successfully disseminated to transmission-relevant tissues despite contrasting early gut dynamics. Furthermore, these strain-specific colonisation dynamics were associated with increased Semliki Forest virus (SFV) transmission and enhanced adult longevity, particularly following colonisation by the exogenous strain. Our findings indicate that resident microbial communities act as selective ecological filters that shape symbiont colonisation in a strain-dependent manner. More broadly, this study demonstrates that the biological effects of mosquito symbionts emerge from interactions between bacterial strains and the microbial community context, providing an ecological framework for the rational and effective development of microbiota-based vector control strategies.

**Importance:** Bacterial symbionts are increasingly being explored as tools to reduce mosquito-borne disease; however, their effects can vary depending on the mosquito host and its microbial community. Thus, a deeper understanding of the interactions among symbionts, their host, and resident microbiota is needed. In this work, we show that resident bacteria can determine whether different *Asaia* strains successfully establish in the invasive mosquito *Aedes koreicus*, demonstrating that symbiont establishment is shaped by interactions within the microbial community rather than by the introduced bacterium alone. Furthermore, successful colonisation was associated with changes in virus transmission and mosquito longevity. These findings show that the microbial environment of a mosquito is an important factor to consider when developing symbiont-based approaches for controlling mosquito-borne diseases.

## Introduction

The discovery that bacterial symbionts can alter arbovirus transmission in mosquito hosts has fundamentally changed our understanding of mosquito biology and established symbionts as a promising target for sustainable vector control^1,2^. For decades, this concept has been largely driven by *Wolbachia*, a maternally inherited intracellular symbiont, whose ability to reduce the transmission of major arboviruses in *Aedes* mosquitoes has led to successful field implementation worldwide to control dengue virus^3–6^. As an intracellular symbiont, *Wolbachia* is now well established to reduce viral replication primarily through host cell-dependent mechanisms, including perturbation of cellular pathways and competition for intracellular resources^7,8^. However, more recently, the discovery that extracellular members of the mosquito microbiota, such as the *Rosenbergiella* strain Y4.6, can also suppress arbovirus infection has broadened this paradigm^9^. Unlike *Wolbachia*, *Rosenbergiella* acts indirectly by modifying the gut microenvironment, reducing gut pH and impairing viral infectivity before cell invasion^9^. Together, these findings demonstrate that microbial inhibition of arboviruses can arise through fundamentally different mechanisms and suggest that mosquito-associated bacteria represent a far more diverse reservoir of antiviral functions than previously recognized.

This expanding repertoire of symbiont-mediated antiviral mechanisms has broadened interest in exploiting bacterial symbionts for disease control while simultaneously exposing a fundamental gap in our understanding: how predictable are symbiont phenotypes across different biological contexts? A major obstacle to answering this question is that microbial symbionts are often treated as functionally homogeneous entities. Many studies have compared the extent to which strain-level variation within a given bacterial taxon functionally shapes colonisation, and host phenotype remains comparatively unaddressed. However, work on both *Wolbachia* and *Rosenbergiella* has demonstrated that strain identity is a key determinant of symbiont biology, influencing bacterial density, tissue tropism, persistence, host interactions, and pathogen interference^10–12^. These findings suggest that symbiont-mediated phenotypes are unlikely to be intrinsic properties of a bacterial species, but instead emerge from interactions among bacterial genotypes, host background, and the resident microbiota^13^. Despite this growing recognition, whether endogenous (native) and exogenous (non-native) strains of the same mosquito symbiont differ in their ability to establish, persist, and influence host biology remains largely unexplored.

To address this question, a model system is needed in which bacterial strain identity can be disentangled from host and microbiota effects: *Asaia* bacteria provide such an opportunity. This acetic acid bacterium is a widespread member of mosquito bacterial communities, naturally colonizing multiple tissues and being efficiently transmitted both vertically and horizontally^14–16^. These features, together with its suitability for genetic manipulation, have made *Asaia* an interesting candidate for microbiota-based vector control through paratransgenesis^17^.

Interestingly, comparative genomic and phylogenomic studies have shown that *Asaia* isolates are not functionally or evolutionarily homogeneous: analyses revealed substantial genomic variation among isolates, including independent signatures of genome reduction and distinct associations with insect hosts^18,19^. These findings suggest that strain identity and host-association history may influence the ecological behaviour of *Asaia* and should therefore be considered when evaluating its colonisation and potential application in vector control.

Despite the translational interest, our understanding of *Asaia*-hosts interactions remains surprisingly limited. Research has focused predominantly on *Anopheles* mosquitoes, where *Asaia* has been explored as a vehicle for delivering anti-*Plasmodium* effectors, leaving the biology of wild-type *Asaia* poorly characterised^17,20,21^. Evidence from *Anopheles* indicates that the effects of wild-type *Asaia* are highly variable: depending on the experimental system, *Asaia* has been associated with immune activation, enhanced *Plasmodium* development through changes in gut physiology, or no relationship with parasite prevalence in field populations^19,22,23^. At the same time, the role in arbovirus systems is limited to a few studies, reporting associations between *Asaia* and altered West Nile virus infection in *Culex* mosquitoes, although bacterial persistence itself appears highly variable^24^. Such contrasting observations suggest that *Asaia*-mediated phenotypes are unlikely to be universal and instead may depend on the ecological context in which the symbiosis occurs.

A potential explanation for these inconsistencies is that *Asaia* has largely been investigated without considering strain identity. Experimental studies commonly employ environmental strains or some originally isolated from different mosquito species, implicitly assuming that they recapitulate the behaviour of naturally occurring host-associated populations. However, endogenous *Asaia* strains have likely co-evolved with their mosquito hosts and exist within a resident microbial community that shapes their establishment, tissue dissemination, and persistence. Whether host-adapted (endogenous) and non-native (exogenous) *Asaia* strains differ in their colonisation dynamics and interactions with the host microbiota therefore remains fundamental, yet largely unexplored.

Recent surveys of the bacterial communities associated with the invasive mosquito species *Aedes koreicus* have identified *Asaia* as a dominant member of its microbiota throughout development and adulthood^25–27^. Moreover, although not considered a primary arbovirus vector, we have demonstrated that *Ae. koreicus* shows competence for several alphaviruses and flaviviruses, making it a suitable model for investigating how host–symbiont interactions may influence vector competence^28,29^.

In this study, we used *Ae. koreicus* to investigate whether symbiont strain origin contributes to the establishment of *Asaia* within the mosquito host. We compared a native *Asaia* isolate from *Ae. koreicus* with the widely used *Anopheles stephensi* strain to determine whether endogenous and exogenous strains differ in their colonisation dynamics and interactions with the resident microbiota. We further assessed whether differences in symbiont colonisation are associated with changes in susceptibility to arbovirus infection. By examining strain identity within its natural host context, we aim to explore how mosquito symbiont ecology informs the development of microbiota-based vector control strategies.

## Results

### Endogenous *Asaia* is dynamically structured across host development and environmental conditions

To define the natural colonisation background against which strain-specific recolonisation could be evaluated, we first characterised endogenous wild-type *Asaia* in *Ae. koreicus*. Under standard insectary conditions, bacterial density changed significantly across development and increased with adult age in both females and males, reaching the highest levels at 20dpe (Figure 1A; females, p = 0.0018; males, p < 0.001, Kruskal–Wallis test). Endogenous *Asaia* was detected in the midgut, salivary glands and reproductive tissues of both sexes, although abundance differed among tissue compartments. In sugar-fed females, bacterial density was higher in the salivary glands than in the ovaries (Figure 1B; p < 0.0001, Kruskal-Wallis). Following blood feeding, *Asaia* remained detectable in the midgut and ovaries, with lower abundance in the ovaries (Figure 1B; p=0.04, Mann-Whitney test). In males, bacterial density was higher in the midgut than in the testes (Figure 1B; p=0.008, Mann-Whitney test).

**Figure 1.**
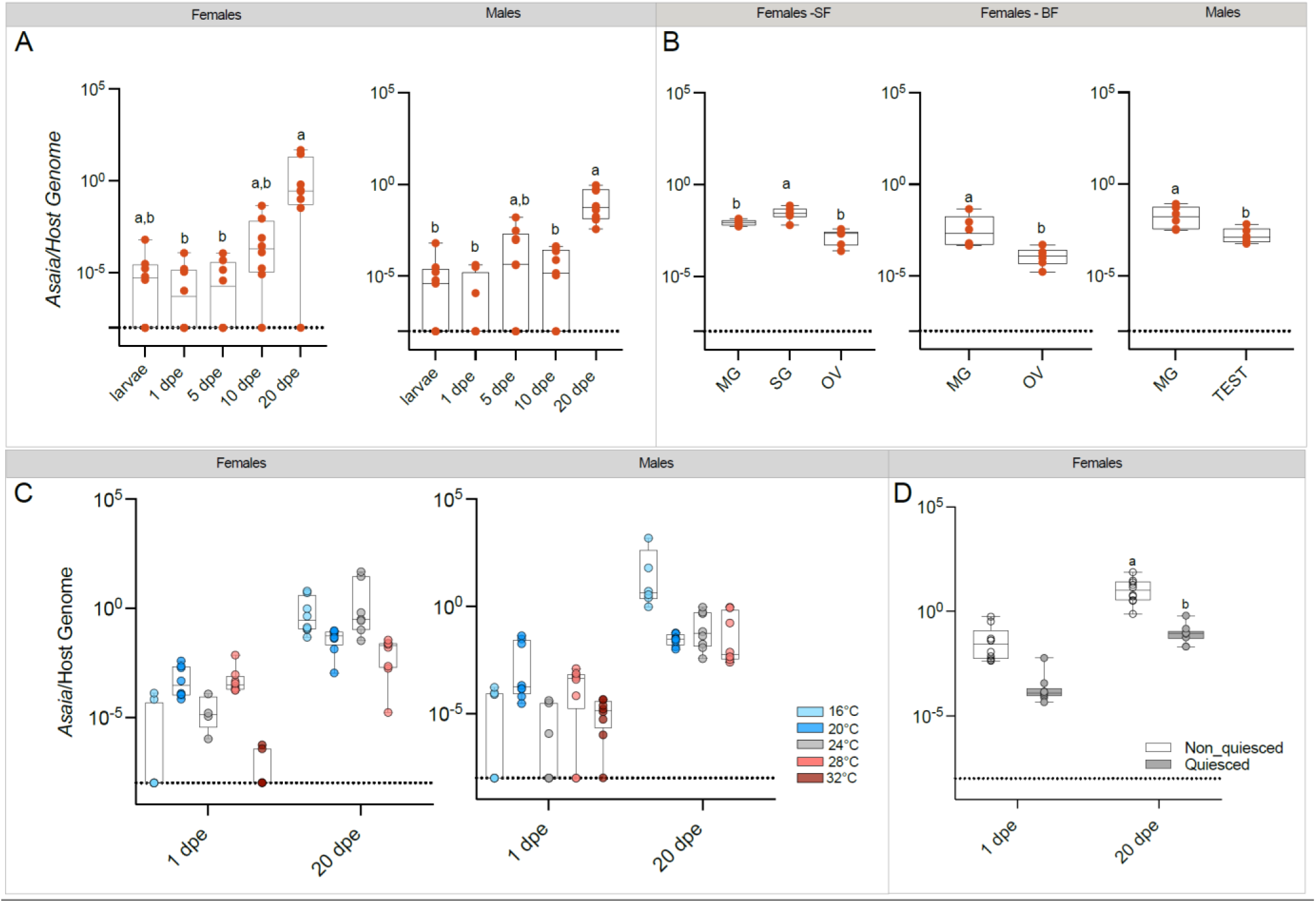
Dynamics and persistence of *Asaia* in *Ae. koreicus*. **(A)** *Asaia* abundance in female and male mosquitoes at different developmental stages. **(B)** *Asaia* abundance in different tissues of sugar-fed females (females-SF), blood-fed females (Females-BF), and male mosquitoes, including midgut (MD), salivary gland (SG), ovaries (OV), and testes (TEST). **(C)** *Asaia* abundance in females and males reared at different temperatures and sampled at 1 and 20 days post-emergence (dpe). **(D)** *Asaia* abundance in females originating from non-quiesced and quiesced eggs at 1 and 20 DPE. In panels A, C, and D, each point represents an individual mosquito, whereas in panel B, each point represents a pool of three tissues from an individual mosquito. Boxplots indicate the distribution of *Asaia* abundance within each group, and the dashed line indicates the limit of detection for *Asaia*. Different letters indicate statistically significant differences between groups, except in panel C. Statistical comparisons are reported in Table S1.

We then assessed whether this endogenous background was affected by environmental and life-history conditions (Figure 1C). Across 138 mosquitoes sampled at various temperatures ranging from 16-32°C, natural *Asaia* density increased significantly from 1 to 20 dpe at every temperature for which both time points were available (16-28°C; all p<0.001). 20 dpe samples were not obtained at 32°C due to heat-associated mortality. By 20dpe, detection of *Asaia* was 100% at every temperature, regardless of detection rate at 1dpe, which ranged from 33-100% depending on the temperature. Natural *Asaia* density was affected by an interaction between temperature and sex (p<0.0001, likelihood-ratio test), indicating that the temperature-density relationship differed between males and females. In females, density was highest at 20°C and 28°C at 1dpe, shifting to a peak at 24°C by 20dpe. In males, density was highest at 20°C at 1dpe, and more broadly elevated across 16-, 24-, and 28°C by 20dpe. Both sexes showed significant temperature-dependent variation in density at both ages (full pairwise comparisons in Supplementary Table S1).

Egg quiescence, a key dormancy trait of *Aedes* mosquitoes, also altered the endogenous *Asaia* profile. Females emerging from quiescent eggs carried lower bacterial densities than those derived from non-quiescent eggs (Figure 1D). Although abundance increased with adult age in both groups, it remained lower in mosquitoes originating from quiescent eggs (quiescence effect, p = 0.0156, F(1,36)). Together, these data establish endogenous *Asaia* as a widespread but dynamic component of the *Ae. koreicus* microbiota, whose abundance varies with adult age, tissue compartment, temperature and egg-quiescence history.

### Host origin shapes *Asaia* colonisation dynamics in *Ae. koreicus*

We observed marked isolate-specific differences in *Asaia* colonisation efficiency and dynamics in *Ae. koreicus,* in both sexes. In both females and males, the proportion of individuals with GFP-positive guts was significantly higher in those colonised with *Asaia ste^gfp^*than in those colonised with *Asaia kor^gfp^* (p = 0.021 and p = 0.005, respectively) (Figure 2 A,B). However there was no significant strain × sex interaction (χ²(1) = 0.66, p = 0.416). Recolonisation prevalence of *Asaia kor^gfp^* increased in both sex from 4 to 12 dpc, and colonisation prevalence differed significantly across time points post-colonisation overall (F(3,25) = 3.18, p=0.041), although no individual comparisons survived correction for multiple testing (Holm-adjusted p≥ 0.10).

**Figure 2.**
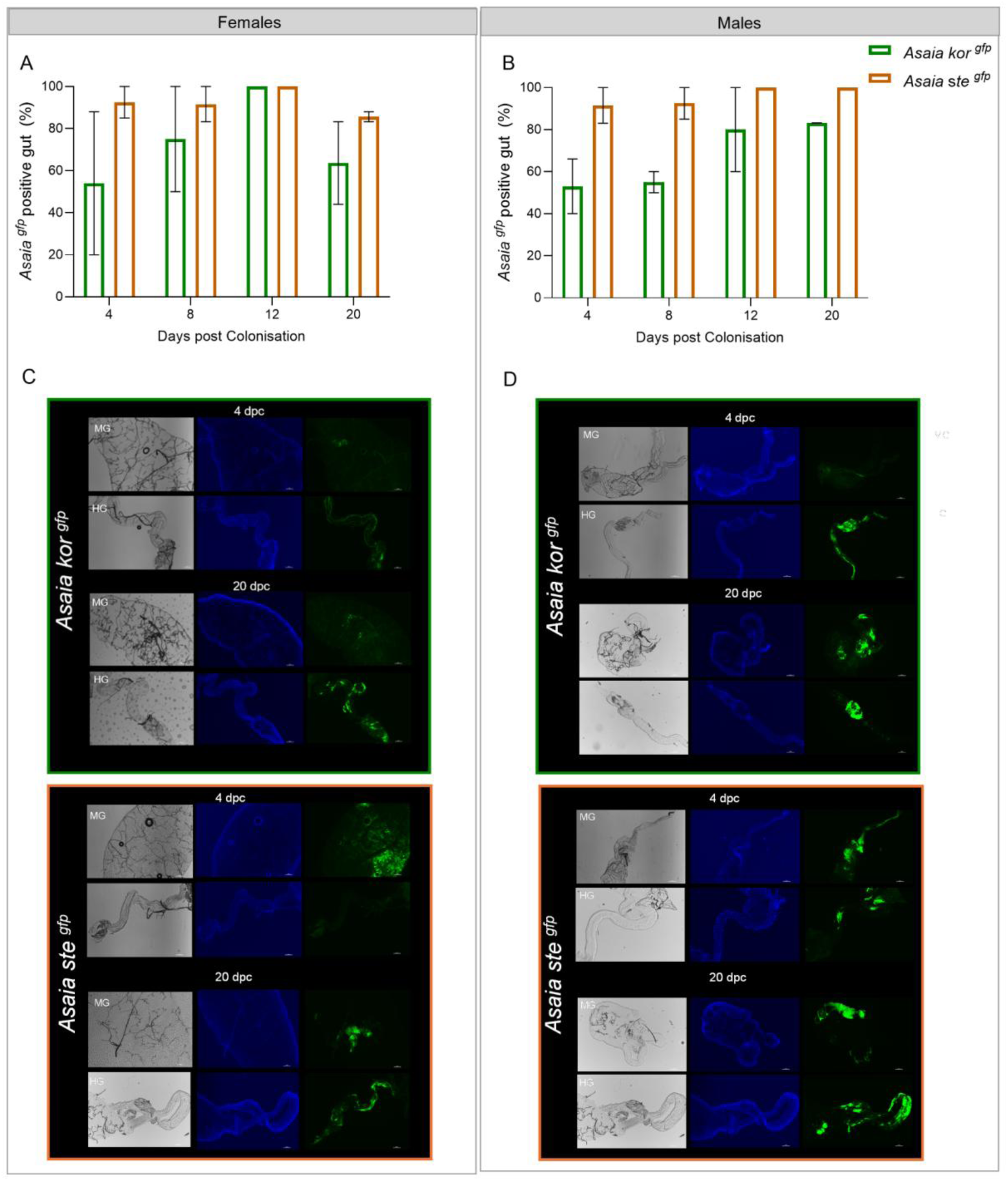
*Asaia* recolonisation dynamics in *Ae. koreicus*. **(A)** Prevalence of *Asaia*-positive females over time post-colonisation. **(B)** Prevalence of *Asaia*-positive males over time post-colonisation. **(C)** Representative fluorescence microscopy images of *Asaia* colonisation in the female midgut (MG) and hindgut (HG). **(D)** Representative fluorescence microscopy images of *Asaia* colonisation in male midgut (MG) and hindgut (HG). For each condition, brightfield, nuclear staining (blue), and GFP fluorescence (green) channels are shown.

Consistent with these differences in prevalence, microscopy revealed distinct strain-specific spatial colonisation patterns within the gut. At 4 dpc, *Asaia ste^gfp^* showed a stronger fluorescent signal and broader distribution across both the midgut and hindgut of males and females compared with *Asaia kor^gfp^*, which appeared largely confined to the hindgut and was not stably detected in the midgut. By 20 dpc, however, the spatial patterns of colonisation appeared more comparable between the two strains in both sexes (Figure 2C,D) (full pairwise comparisons in Supplementary Table S2 and S3).

### The microbial composition shapes colonisation of the endogenous *Asaia* strain

To determine whether strain-specific *Asaia* colonisation reflects intrinsic bacterial properties or ecological filtering by the resident microbiota, we established an axenic (germ-free) *Ae. koreicus* platform that allowed the host microbial environment to be experimentally controlled without detectable effects on mosquito fitness (Figure S1). This system enabled us to disentangle the contribution of bacterial strain identity from that of the native microbial community by comparing recolonisation by the endogenous *Asaia kor^gfp^* and exogenous *Asaia ste^gfp^* isolates in wild type (WT) and axenic (AX) hosts, assessed separately in gut and carcass, at 4, 8, and 12 days post-colonisation (dpc; 150 mosquitoes per compartment).

In the gut (Figure 3B-C, prevalence; Figure 3F-G, bacterial load), colonisation prevalence was high by 4dpc for *Asaia kor^gfp^*AX, *Asaia ste^gfp^* WT and *Asaia ste^gfp^* AX (92-100%), whereas *Asaia kor^gfp^* WT colonised more slowly (42% at 4dpc); all four combinations reached 100% prevalence by 12dpc. Microbiota status also significantly affected bacterial load in a strain-dependent manner (strain x condition interaction, p<0.0001). At 4dpc, *Asaia kor^gfp^* reached a far lower bacterial load when compared to *Asaia ste^gfp^* in WT guts (∼7,000-fold, p<0.0001), but this gap narrowed substantially in axenic guts (∼11-fold, p=0.012), indicating that removal of the native microbiota partially rescues early *Asaia kor^gfp^* proliferation. This rescue was transient, reversing by 8dpc, and converging by 12dpc.

**Figure 3.**
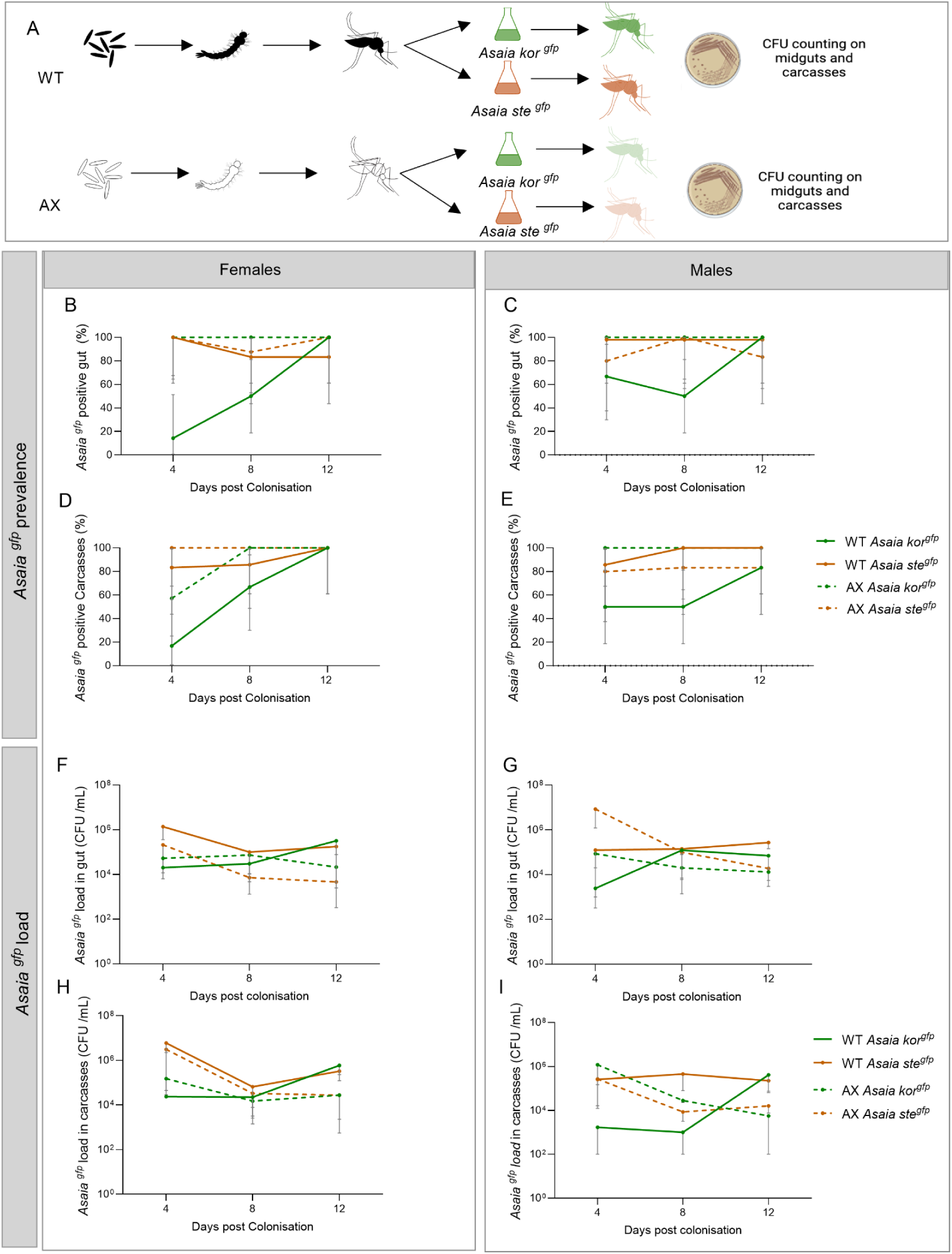
Comparison of *Asaia* recolonisation dynamics in wild-type and axenic *Ae. koreicus*. (A) Experimental design. (B) Prevalence (%) of *Asaia ^gfp^*-positive gut samples in females. (B) Prevalence (%) of *Asaia ^gfp^*-positive gut samples in males. (C) Prevalence (%) of *Asaia ^gfp^*-positive carcass samples in females. (D) Prevalence (%) of *Asaia ^gfp^*-positive carcass samples in males. (E) *Asaia ^gfp^* load in gut samples of females. (F) *Asaia ^gfp^* load in gut samples of males. (G) *Asaia ^gfp^* load in carcass samples of females. (H) *Asaia ^gfp^* load in carcass samples of males. Points represent mean values ± SD, Statistical comparisons are reported in Tables S2 and S3.

A similar prevalence pattern was seen in the carcass (Figure 3D-E, prevalence; Figure 3H-I, bacterial load), with *Asaia ste^gfp^* AX and WT being highly prevalent by 4dpc (92%), and *Asaia kor^gfp^*AX reaching 100% prevalence by 8dpc, but *Asaia kor^gfp^* WT again lagged, reaching only 33% prevalence at 4dpc, before rising to 92% by 8 and 12dpc. However, unlike prevalence, bacterial load showed no equivalent rescue effect. *Asaia kor^gfp^* loads did not differ by microbiota status at 4dpc (p=0.22), and the strain x condition interaction was not significant overall (p=0.99). From 8dpc onwards, *Asaia ste^gfp^*reached a higher load than *Asaia kor^gfp^* in both compartments regardless of condition (p<0.04). Sex had a significant additive effect on load in the carcass only (males ∼2.3-fold lower than females, p=0.035), with no sex interactions detected in either compartment; statistical comparisons of strain, condition and timepoint are therefore reported pooled across sex.

### Host-origin-dependent *Asaia* colonisation alters SFV transmission in *Ae. koreicus*

To determine whether strain-specific *Asaia* colonisation influences arbovirus infection in *Ae. koreicus*, we assessed Semliki Forest virus (SFV) infection and transmission following re-colonisation with the endogenous *Asaia kor* and exogenous *Asaia ste* strains. Because salivary gland colonisation is particularly relevant to viral dissemination and transmission, we first quantified bacterial loads of *Asaia kor* and *Asaia ste* in the salivary glands before and after a blood meal. The abundance of both strains remained stable following blood feeding, and both displayed consistent colonisation of the salivary glands (Figure S2).

We next examined whether recolonisation altered SFV infection and dissemination. The infection rate was higher in mosquitoes recolonised with *Asaia ste* (62.5%) than in control mosquitoes (41.7%), whereas females recolonised with *Asaia kor* showed a slightly lower infection rate (39.1%) relative to controls (Figure 4A). Nevertheless, a trend towards higher carcass viral loads was observed in mosquitoes recolonised with both *Asaia* strains compared with controls; however, these differences were not statistically significant (Figure 4C).

**Figure 4.**
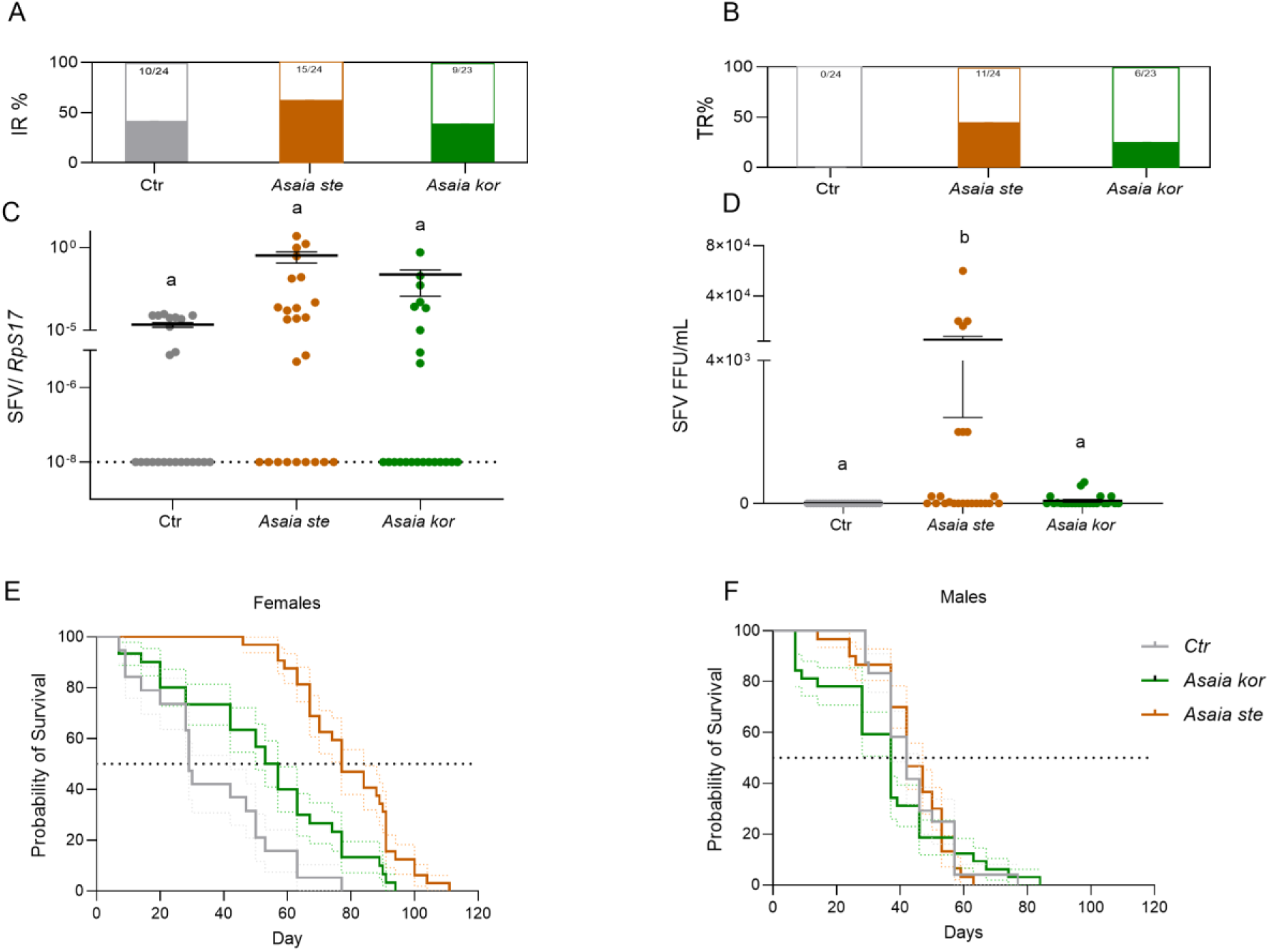
*Asaia* effect on SFV infection in *Ae. koreicus* and its effect on adult survival. **(A)** SFV IR% (infection rate) in mosquito carcasses following exposure to *Asaia*. **(B)** SFV TR% (transmission rate) in salivary glands following exposure to *Asaia*. **(C)** Viral load in mosquito carcasses following exposure to *Asaia*. **(D)** Viral load in mosquito salivary glands following exposure to *Asaia*. **(E)** Survival curves of female mosquitoes following an 8-day exposure to *Asaia kor ^gfp^* or *Asaia ste ^gfp^* compared with the control (CTR) group. **(F)** Survival curves of male mosquitoes following an 8-day exposure to *Asaia kor ^gfp^* or *Asaia ste ^gfp^*compared with the control (CTR) group. In C and D, each dot represents an individual mosquito. Error bars indicate mean ± SEM. Dashed lines indicate the limit of detection where applicable.

The effect of *Asaia* recolonisation became more pronounced at the transmission stage. No transmission was detected in control mosquitoes, whereas transmission rates reached 26.0% in females recolonised with *Asaia kor* and 45.8% in those recolonised with *Asaia ste* (Figure 4B). Consistent with this pattern, salivary gland viral titres were significantly higher in mosquitoes recolonised with *Asaia ste* than in controls (p=0.0004, Kruskal-Wallis test) (Figure 4D). Although low levels of virus were also detected in the salivary glands of mosquitoes carrying *Asaia kor*, viral loads did not differ significantly from controls. No significant difference was detected between the two recolonised groups.

Because adult survival is a key life-history trait contributing to vectorial capacity, we next assessed whether *Asaia* supplementation affected female longevity. Females recolonised with *Asaia* survived significantly longer than control mosquitoes (p<0.0001, Logrank Mantel-Cox test, Figure 4F,G), indicating that bacterial supplementation influenced host survival in addition to its association with SFV transmission.

## Discussion

Interest in microbiota-based approaches to mosquito control has broadened research into bacterial symbionts capable of modifying host biology or pathogen transmission^2^. Our findings indicate that predicting symbiont function requires moving beyond the bacterium itself and considering the ecological composition in which it operates. *Asaia* colonisation in *Ae. koreicus* emerged as the product of interactions between the bacterial strain, the host and its resident microbial community. Using endogenous and exogenous *Asaia* strains as an experimental framework, we showed that these ecological interactions shape bacterial establishment within the host and are associated with differences in arbovirus transmission capacity.

*Ae. koreicus* proved to be a particularly informative system for investigating these interactions, as its association with *Asaia* appears to differ from that observed in *Ae. albopictus* and *Ae. aegypti*, where the bacterium is generally detected at lower relative abundance^25,30^. Studies on field and laboratory samples showed that *Ae. koreicus* naturally harbours abundant *Asaia* populations throughout development and across multiple tissues^27^. Our baseline characterisation further demonstrated that this endogenous population is highly dynamic: although *Asaia* was consistently detected throughout the mosquito life cycle, its abundance varied with adult age, tissue compartment, temperature and egg-quiescence history. These observations indicate that even the naturally associated symbiont is continuously shaped by host development and environmental conditions, emphasizing that microbial abundance is itself a dynamic ecological trait.

Within this ecological framework, bacterial strain identity represents one component contributing to colonisation outcomes. The endogenous (*Asaia kor^gfp^*) and exogenous *(Asaia ste^gfp^*) isolates exhibited distinct colonisation dynamics during the early stages of host establishment (4-8 days post-colonisation). The exogenous strain colonised the gut more rapidly and disseminated earlier, whereas the endogenous strain initially remained spatially restricted before progressively reaching comparable tissue distributions. These findings suggest that different *Asaia* strains interact distinctly with the host environment, although these effects are neither absolute nor consistent across tissues. Instead, colonisation trajectories depend on both bacterial identity and the biological context in which colonisation occurs. Although intrinsic differences between the *Asaia* isolates may contribute to their contrasting behaviour ^18^, our experimental design indicates that colonisation is also shaped by the host ecological context and its resident microbiota.

The axenic recolonisation experiments further illustrate the importance of this context. By experimentally removing the resident microbiota while maintaining mosquito fitness, we were able to disentangle bacterial properties from ecological interactions occurring within the host. Recent ecological frameworks propose that host-associated microbiota should be viewed as dynamic communities assembled through processes such as environmental filtering, community interactions, and priority effects rather than collections of independently acting microorganisms^31^. Within this framework, our findings suggest that the resident microbiota of *Ae. koreicus* functions as a selective ecological filter, constraining the establishment of the endogenous *Asaia* strain while exerting comparatively little influence on the exogenous isolate we have tested.

This pattern suggests that the native isolate is more tightly integrated into the homeostatic mechanisms regulating the resident microbial community. In established host-microbiota associations, long-term persistence does not necessarily require maximal bacterial growth and spread, but rather the maintenance of stable population sizes through interactions involving the host, neighbouring microorganisms and immune regulation^32,33^. The endogenous *Asaia* population may therefore be subject to ecological constraints that limit its early expansion while ensuring long-term persistence within the host. In contrast, the exogenous strain appeared comparatively less affected by these constraints, suggesting that it interacts differently with the resident microbial community, potentially due to the occupation of alternative ecological niches or experiencing different competitive interactions.

Furthermore, these distinct colonisation patterns were associated with differences in arbovirus transmission. Both *Asaia* strains successfully colonised the salivary glands and remained stable following blood feeding, indicating that early differences in gut colonisation do not necessarily predict dissemination to transmission-relevant tissues. Nevertheless, recolonisation with the exogenous *Asaia* strain was associated with the highest SFV transmission rate and significantly elevated viral titres in the salivary glands. Previous work in malaria mosquitoes has shown that *Asaia* can influence mosquito physiology by altering glucose metabolism and gut pH, thereby promoting *Plasmodium* development^23^. Whether similar indirect physiological effects contribute to arbovirus susceptibility needs further investigation.

Interestingly, the consequences of *Asaia* supplementation extended beyond viral infection. Females supplemented with *Asaia* survived significantly longer than control mosquitoes. Because adult longevity is a major determinant of vectorial capacity, increasing the probability that mosquitoes survive the extrinsic incubation period and remain capable of transmitting infection, this result suggests that *Asaia* may influence multiple components of transmission potential. Previous studies have also reported positive effects of *Asaia* on larval survival and development, possibly through contributions to host nutrition and metabolism^24,34–36^. Although the mechanisms underlying the increased adult longevity observed here remain to be determined, these findings reinforce the view that the consequences of symbiosis should be evaluated at the level of the whole-host phenotype rather than solely through direct effects on pathogen infection or blocking.

Collectively, our findings suggest that evaluating candidate symbionts solely according to bacterial species provides an incomplete understanding of their biology and their potential for vector control. Instead, symbiont performance emerges from the interaction between bacterial strain identity, host background and the resident microbial community. As microbiota-based strategies continue to expand, incorporating this ecological complexity will be essential for selecting, engineering and deploying bacterial symbionts with predictable outcomes.

## Materials and methods

### Mosquitos rearing

A colony of *Ae. koreicus* was established from larvae collected in Belluno Province, Veneto Region, Italy (46.0139823 N, 11.8971688 E), and has been maintained in the insectary of the University of Pavia since 2023, as described in Bahrami et al., 2026 ^29^. Mosquitoes were reared at 24 ± 0.5°C, 70% relative humidity, and a 14:10 h light photoperiod. Larvae were maintained in dechlorinated water and fed TetraMin flake fish food. Adults were maintained in mesh cages (BugDorm; 60 × 60 × 120 cm) with 10% sucrose *ad libitum*. Females were blood-fed using a membrane feeding system (Hemotek, UK) with human blood obtained from Centro Trasfusionale, IRCCS Policlinico San Matteo, Pavia, Italy. Oviposition cups were provided 7–10 days post-blood feeding.

### *Asaia* Strains and Culture Conditions

Two GFP-expressing *Asaia* strains were used in this study, *Asaia* sp. SF2.1(Gfp), originally isolated from an adult female *Anopheles stephensi* and previously described by Favia et al. (2007)^15^, is hereafter referred to as *Asaia ste^gfp^*. *Asaia* sp. AkF3, originally isolated from *Aedes koreicus* and described by Comandatore et al. (2021)^18^, was transformed with the GFP-expressing plasmid pHM2-Gfp for the present study and is hereafter referred to as *Asaia kor^gfp^*. Transformation of *Asaia kor^gfp^* was performed by electroporation as previously described by Favia et al. (2007)^15^. For the experiments, both GFP-expressing strains were cultured for 24 h at 30 °C in GLY medium (25 mL/L glycerol, 10 g/L yeast extract; pH 5.0). Cultures were grown to an optical density at 600 nm (OD600) of 1.0 (approximately 1 × 10⁸ cells/mL), harvested by centrifugation, washed three times in a physiological solution containing 0.9% NaCl, and resuspended in sterile 5% (w/v) sucrose solution.

### Quantification of *Asaia* density in *Ae. koreicus* under standard and stress conditions

#### Baseline *Asaia* abundance and tissue distribution

To characterize baseline *Asaia* density in the laboratory colony of *Ae. koreicus*, mosquitoes were reared under standard rearing conditions. Virgin females and males were collected at 1, 5, 10, and 20 days post-emergence (dpe). At each time point, eight individuals of each sex and eight larvae were collected. To assess tissue-specific *Asaia* distribution, sugar-fed virgin females and males were collected at 9 dpe. Following surface sterilization ^37^, midguts, salivary glands, and ovaries were dissected from females, whereas midguts and testes were dissected from males. For each sex, three tissues were pooled per biological replicate, with six biological replicates per condition. In a separate cohort, females were blood-fed at 8 dpe and 18 blood-fed individuals were collected 24 h later. Ovaries and midguts were dissected and pooled into six replicates.

#### Effect of egg quiescence and temperature on *Asaia* abundance

To assess the effects of abiotic stress, eggs were maintained under quiescent conditions for 11 weeks before hatching. Resulting females were reared under standard conditions and collected at 1 and 20 dpe. Controls were derived from eggs quiesced for 4 weeks. For temperature treatments, mosquitoes were reared continuously at 16, 20, 24, 28, or 32°C. Females and males were collected at 1 and 20 dpe, with eight individuals of each sex analysed per condition. At 32°C, mortality prevented collection at 20 dpe.

#### DNA extraction and qPCR

Genomic DNA was extracted using the Wizard® Genomic DNA Purification Kit (Promega Corporation, Madison, WI, USA) according to the manufacturer’s instructions and eluted in 20 μL of DNase-free water. DNA concentration was measured using a NanoDrop ND-1000 spectrophotometer (Thermo Fisher Scientific, Wilmington, DE, USA). *Asaia* abundance was quantified by quantitative PCR (qPCR) using *Asaia*-specific primers, with the mosquito *homothorax* gene (AALC636_001297) used as the reference gene(Table S4). qPCR reactions were performed in a final volume of 10 μL containing 5 μL of 2× QuantiNova SYBR Green Master Mix (QIAGEN, Hilden, Germany), 0.5 μL each of forward and reverse primers, 2 μL of nuclease-free water, and 2 μL of template DNA. Reactions were performed at 95°C for 2 min, followed by 40 cycles of 95°C for 5 s and 60°C for 30 s, with a subsequent melt-curve analysis.

### Generation of axenic mosquitoes and validation

#### Axenic mosquito generation

Germ-free mosquitoes (hereafter axenic mosquitoes) were generated by adapting previously described protocols^38,39^. All procedures were performed under sterile conditions inside a microbiological safety cabinet. Eggs were surface sterilised by sequential immersion in 70% ethanol (5 min), 1% sodium hypochlorite (5 min), and 70% ethanol (5 min), with sterile water rinses between steps. Approximately 100 eggs were maintained in sterile water per replicate (n = 4 replicates per experiment).

Axenic larvae (L1) were reared in sterile conditions and temporarily colonised with *Escherichia coli* (2–5 × 10⁸ CFU/mL) supplemented with autoclaved TetraMin Baby food. At the L4 stage, ampicillin (100 μg/mL) was applied to eliminate the provided *E. coli*, and larvae were maintained under sterile conditions until pupation. Pupae were surface-sterilised and allowed to emerge in sterile containers.

#### Sterility validation and developmental assessment

Sterility was assessed at the egg, larval, and adult stages. For egg validation, subsets of 20–30 surface-sterilised eggs from each experimental group were inoculated into Luria-Bertani (LB) medium and monitored for microbial growth. At the larval stage, newly hatched individuals (n = 8 per group) were maintained in sterile water and provided with autoclaved food. Lack of normal development was used as an additional indicator of successful axenic rearing. Additionally, adult mosquitoes were tested for sterility. Individuals (n = 6 per time point) were homogenised without surface sterilisation in 100 μL of sterile 1× PBS and plated onto LB agar and Malt Extract Agar (MEA) to detect residual bacterial and yeast contamination, respectively. Plates were monitored for microbial growth, and the absence of growth on both media was used to confirm axenic status in comparison with wild-type mosquitoes (WT, non-sterilised eggs and reared under standard insectary conditions) and conventional mosquitoes (CN, surface-sterilised eggs and reared subsequently under standard insectary conditions). Axenic status was further assessed by qPCR using bacteria-specific 16S rRNA gene primers (Table S4). Batches showing bacterial growth were excluded from downstream analyses.

To evaluate the effect of axenic conditions on development, larval development time (LDT), pupal development time (PDT), pupation rate, and adult emergence rate were recorded daily under identical environmental conditions across two biological replicates.

### *Asaia* strain-specific re-colonisation

#### *Asaia* (Gfp) re-colonisation

Virgin male and female mosquitoes reared under standard conditions, immediately after emergence, were allowed to feed *ad libitum* for 8 days on 5% sucrose solution supplemented either *Asaia kor^gfp^* or *Asaia ste^gfp^*. Afterwards, mosquitoes were maintained on 5% sucrose supplemented with 100 μg/mL kanamycin. Individuals were sampled at 4, 8, 12, and 21 days post-colonisation (dpc) (8 males and 8 females per group and time point). Following surface sterilisation, midguts were dissected and examined for GFP fluorescence using an Olympus CKX53 inverted microscope. Guts collected at 4 and 20 dpc were fixed in 4% paraformaldehyde for 10 min at 4°C, mounted in glycerol–PBS, and examined by confocal microscopy. Control mosquitoes were maintained without bacterial supplementation. Experiments were performed in two independent biological replicates.

#### Effect of blood feeding on colonization

A subset of virgin females reared under standard condition exposed to either *Asaia kor^gfp^* or *Asaia ste^gfp^* was collected at 8 dpc. To evaluate the effect of blood feeding, females were starved for 24 h and then offered a blood meal. After feeding, mosquitoes were maintained on 5% sucrose containing 100 μg/mL kanamycin. At 1 day and 7 days post-blood meal, 10 individuals per time point and strain were collected. After surface sterilization, salivary glands were dissected from individual mosquitoes and homogenized separately in 100 μL sterile 1× PBS. Homogenates were serially diluted and plated on GLY agar supplemented with kanamycin (100 μg/mL). Plates were incubated at 30°C for 48 h, after which *Asaia* colony-forming units (CFU) were counted manually. Bacterial density was calculated from the number of colonies and the corresponding dilution factor and expressed as CFU/mL.

#### Adult survival

To assess the impact of *Asaia* colonisation on mosquito survival, adult longevity was monitored following exposure for 8 days to either *Asaia kor^gfp^* or *Asaia ste^gfp^*. After colonization, survival was recorded daily until all individuals had died. At least 32 mosquitoes per condition and sex were analysed, alongside control groups (n = 24 per sex).

### Comparative *Asaia re-*colonisation in WT and axenic mosquitoes

Virgin male and female mosquitoes from WT and axenic groups were exposed after emergence to either *Asaia kor^gfp^* or *Asaia ste^gfp^*, as described above. Mosquitoes from each treatment were sampled at 4, 8, and 12 days post-colonisation (dpc), with six individuals per sex, strain, mosquito status, and time point. Following surface sterilisation, midguts and corresponding carcasses were dissected, homogenized and processed for CFU counting as described before.

### Viral challenges

#### SFV infection and mosquito sampling

Females previously colonised with *Asaia ste* or *Asaia kor* for 8 days were exposed to an infectious blood meal containing Semliki Forest virus (SFV-4) at a final concentration of 1 × 10⁸ PFU/mL. At 7 days post-infection (dpi), females were collected and surface-sterilised as described previously. Mosquitoes were dissected for analysis, with salivary glands transferred to Glasgow’s minimum essential medium (GMEM; Gibco, Thermo Fisher Scientific, Waltham, MA, USA) for quantification of virus titre by fluorescent focus assay (FFA), and carcasses transferred to TRIzol® Reagent (Invitrogen, Thermo Fisher Scientific) for dissemination analysis by RT-qPCR.

#### Viral dissemination by RT-qPCR

Individual carcasses were homogenised in 400µL TRIzol® and total RNA was extracted following the manufacturer’s instructions. cDNA synthesis and RT-qPCR were performed as described above using SFV4 E1-specific primers. Samples were considered positive for viral dissemination when amplification was detected within 40 cycles, and expression was normalised to the *Ae. koreicus* ribosomal protein S17 (RpS17) reference gene(Table S4).

#### Infectious virus in salivary glands

Salivary glands were homogenised in GMEM and serially diluted, 50µL of each dilution was added to confluent Baby Hamster Kidney 21 cells (BHK21) in a 96-well plate, and incubated for 1 hour. Dilutions were removed and replaced with an Avicel overlay medium (50:50 mixture of 2% Avicel and GMEM supplemented with 5% foetal bovine serum), and incubated for 24hr at 37°C with 5% CO_2_. Cells were fixed and stained with an alphavirus monoclonal antibody (G77L; Thermo Fisher Scientific) followed by goat anti-mouse Alexa Fluor 488 (A-11001; Thermo Fisher Scientific). Fluorescent foci were counted using an Olympus CKX53 inverted fluorescent microscope and virus titres were expressed as FFU/mL.

### Statistical analysis

Statistical analyses were performed using GraphPad Prism version 8 (GraphPad Software, San Diego, CA, USA) and R version 4.6.1. Data distributions were assessed using the Shapiro–Wilk test^40^. Parametric data were analysed using one- or two-way ANOVA followed by Tukey’s multiple-comparisons test, whereas nonparametric data were analysed using Mann–Whitney *U* or Kruskal–Wallis tests followed by Dunn’s test. Survival was analysed using Kaplan–Meier curves and the log-rank (Mantel–Cox) test.^41^.

*Asaia* colonisation prevalence was modelled in R using a binomial generalised linear model (GLM) with a logit link, with *Asaia* strain, sex and their interaction fitted as fixed effects; days post-colonisation was included as an additive effect; standard errors were adjusted for overdispersion. Post-hoc pairwise comparisons (strain within each sex; between time points) used estimated marginal means (emmeans) with Holm correction.

Natural *Asaia* density and experimental colonisation (gut and carcass compartments) were analysed in R using two-part (“hurdle”) models implemented in the glmmTMB package^42^, combining a Bernoulli component for the probability of detection/colonisation with a conditional magnitude component for bacterial load (Gamma distribution for density; zero-truncated negative binomial for colony forming unit counts). For the experimental challenge assays *Asaia* strain, microbiota condition (wild-type or axenic) and days post-colonisation were fitted as fixed effects with a full three-way interaction; for natural density, temperature and timepoint were fitted with their interaction. Sex was included as an additive covariate as likelihood-ratio tests showed no consistent interaction. Hurdle models were selected over standard count models by Akaike Information Criterion (AIC) and validated using DHARMa residual diagnostics. Post-hoc pairwise comparisons used estimated marginal means (emmeans; Lenth, 2016) with Holm correction. For all analyses, *P* ≤ 0.05 was considered statistically significant.

## Acknowledgments

We thank Sara Repetto and Gianmarco Rotondi for supporting with mosquito colony maintenance and laboratory activities. We also thank Mariangela Bonizzoni for the constructive discussions.

## Data availability

All datasets generated and analysed during this study are publicly available under the DOI 10.5281/zenodo.22674992

---

**Figure S1.**
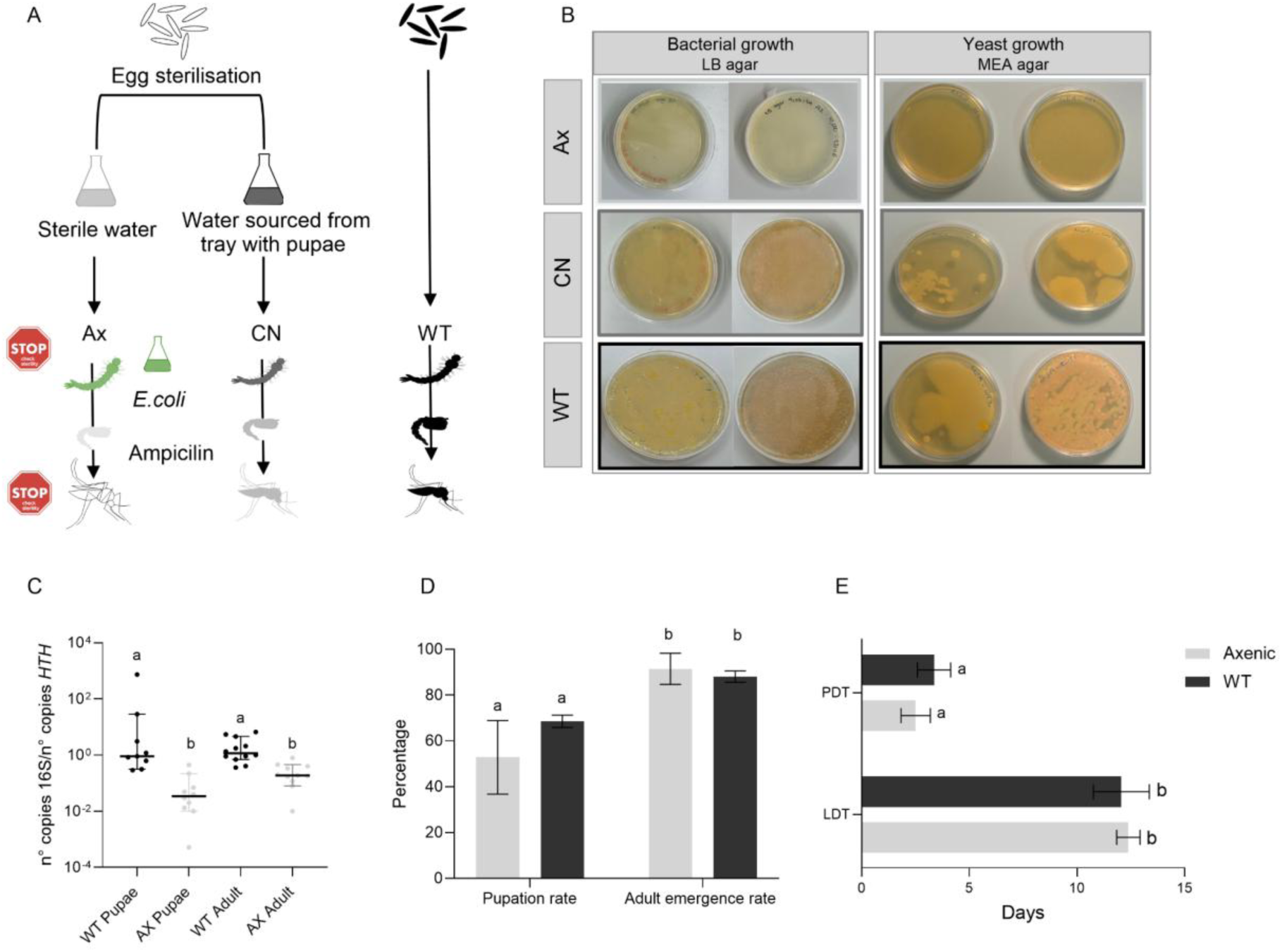
Validation of axenic *Aedes koreicus* mosquitoes and effects of microbiota depletion on development. **(A)** Experimental design to generate axenic (Ax), conventional (CN), and wild-type (WT) mosquitoes. **(B)** Sterility assessment of Ax, CN, and WT mosquitoes by culturing samples on LB agar to detect bacterial growth and MEA agar to detect yeast growth. **(C)** Relative bacterial load quantified by 16S rRNA gene copy number normalised to the mosquito housekeeping gene *HTH* in pupae and adults. Each point represents an independent sample. Horizontal lines indicate the mean ± SD. **(D)** Pupation and adult emergence rates in Ax and WT mosquitoes. **(E)** Larval development time (LDT) and pupal development time (PDT) in Ax and WT mosquitoes. Bars represent mean ± SD.

**Figure S2.**
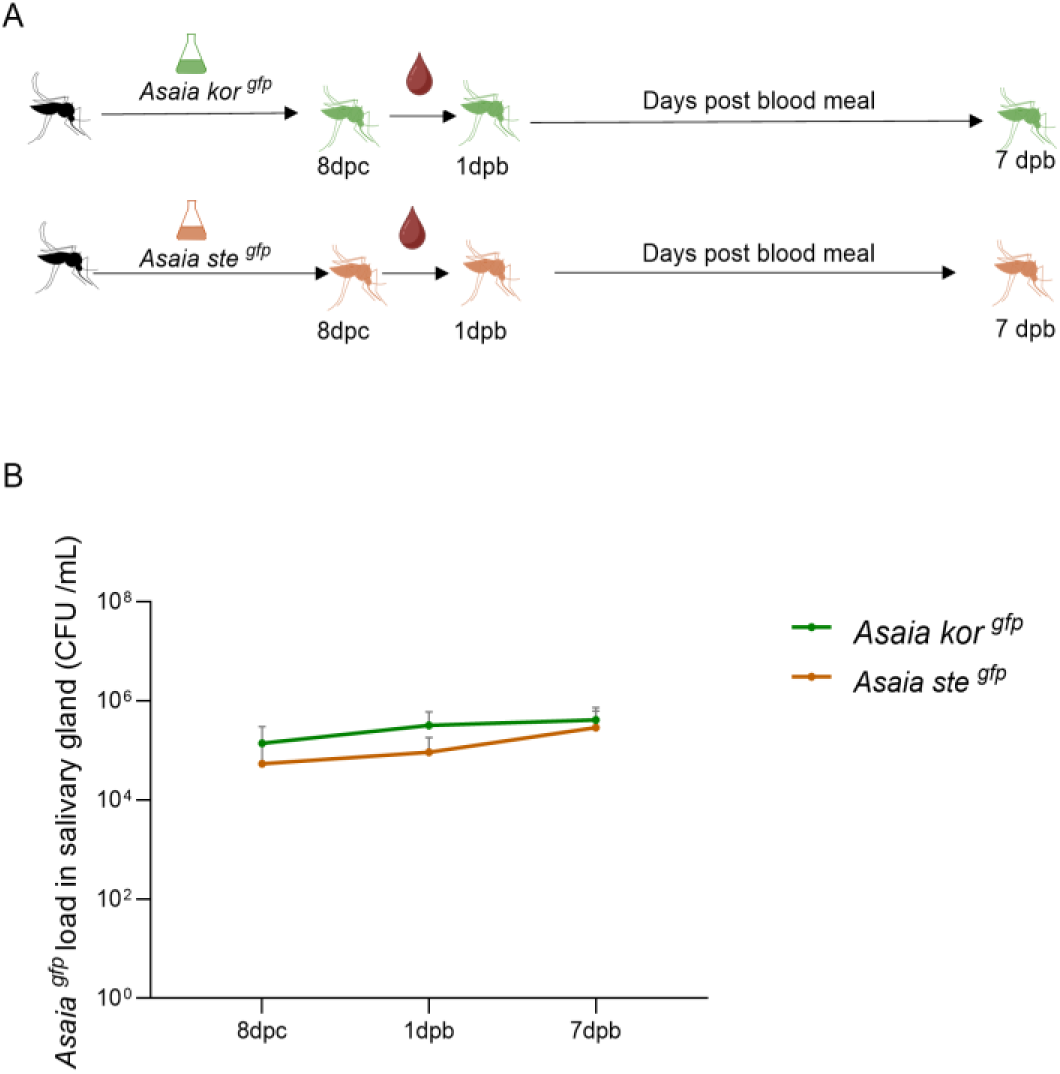
*Asaia* recolonisation dynamics in salivary glands before and after blood meal in Ae.koreicus. **(A)** Experimental design. **(B)** *Asaia ^gfp^* load in salivary gland samples of females after 8 days post-colonisation (dpc) with *Asaia ^gfp^* and following the blood meal, expressed as days post-blood meal (dpb). Points represent mean values ± SD.

## Supplementary

**Table S1.** Pairwise comparisons of natural *Asaia* abundance after thermal stress, by sex and time point.

| Sex | dpe(days post emergence) | Comparison | Ratio | SE(standard error ) | p value |
| --- | --- | --- | --- | --- | --- |
| Female | 1 | 16°C vs 20°C | 0.090 | 0.057 | 0.0004 |
| Female | 1 | 16°C vs 24°C | 2.69 | 1.80 | 0.277 |
| Female | 1 | 16°C vs 28°C | 0.081 | 0.036 | <0.0001 |
| Female | 1 | 16°C vs 32°C | 219 | 141 | <0.0001 |
| Female | 1 | 20°C vs 24°C | 29.9 | 25.6 | 0.0003 |
| Female | 1 | 20°C vs 28°C | 0.90 | 0.63 | 0.882 |
| Female | 1 | 20°C vs 32°C | 2430 | 2040 | <0.0001 |
| Female | 1 | 24°C vs 28°C | 0.030 | 0.022 | <0.0001 |
| Female | 1 | 24°C vs 32°C | 81.3 | 70.5 | <0.0001 |
| Female | 1 | 28°C vs 32°C | 2700 | 1920 | <0.0001 |
| Female | 20 | 16°C vs 20°C | 30.9 | 20.8 | <0.0001 |
| Female | 20 | 16°C vs 24°C | 0.148 | 0.131 | 0.063 |
| Female | 20 | 16°C vs 28°C | 103.6 | 86.9 | <0.0001 |
| Female | 20 | 20°C vs 24°C | 0.0048 | 0.0033 | <0.0001 |
| Female | 20 | 20°C vs 28°C | 3.36 | 2.11 | 0.063 |
| Female | 20 | 24°C vs 28°C | 702 | 600 | <0.0001 |
| Male | 1 | 16°C vs 20°C | 0.0098 | 0.0060 | <0.0001 |
| Male | 1 | 16°C vs 24°C | 4.57 | 3.41 | 0.084 |
| Male | 1 | 16°C vs 28°C | 0.188 | 0.090 | 0.0015 |
| Male | 1 | 16°C vs 32°C | 5.39 | 2.02 | <0.0001 |
| Male | 1 | 20°C vs 24°C | 466 | 432 | <0.0001 |
| Male | 1 | 20°C vs 28°C | 19.2 | 14.0 | 0.0003 |
| Male | 1 | 20°C vs 32°C | 550 | 366 | <0.0001 |
| Male | 1 | 24°C vs 28°C | 0.041 | 0.035 | 0.0006 |
| Male | 1 | 24°C vs 32°C | 1.18 | 0.93 | 0.834 |
| Male | 1 | 28°C vs 32°C | 28.7 | 15.6 | <0.0001 |
| Male | 20 | 16°C vs 20°C | 8475 | 6470 | <0.0001 |
| Male | 20 | 16°C vs 24°C | 1035 | 964 | <0.0001 |
| Male | 20 | 16°C vs 28°C | 1095 | 998 | <0.0001 |
| Male | 20 | 20°C vs 24°C | 0.122 | 0.080 | 0.0034 |
| Male | 20 | 20°C vs 28°C | 0.129 | 0.081 | 0.0034 |
| Male | 20 | 24°C vs 28°C | 1.06 | 0.87 | 0.946 |

**Table S2.**
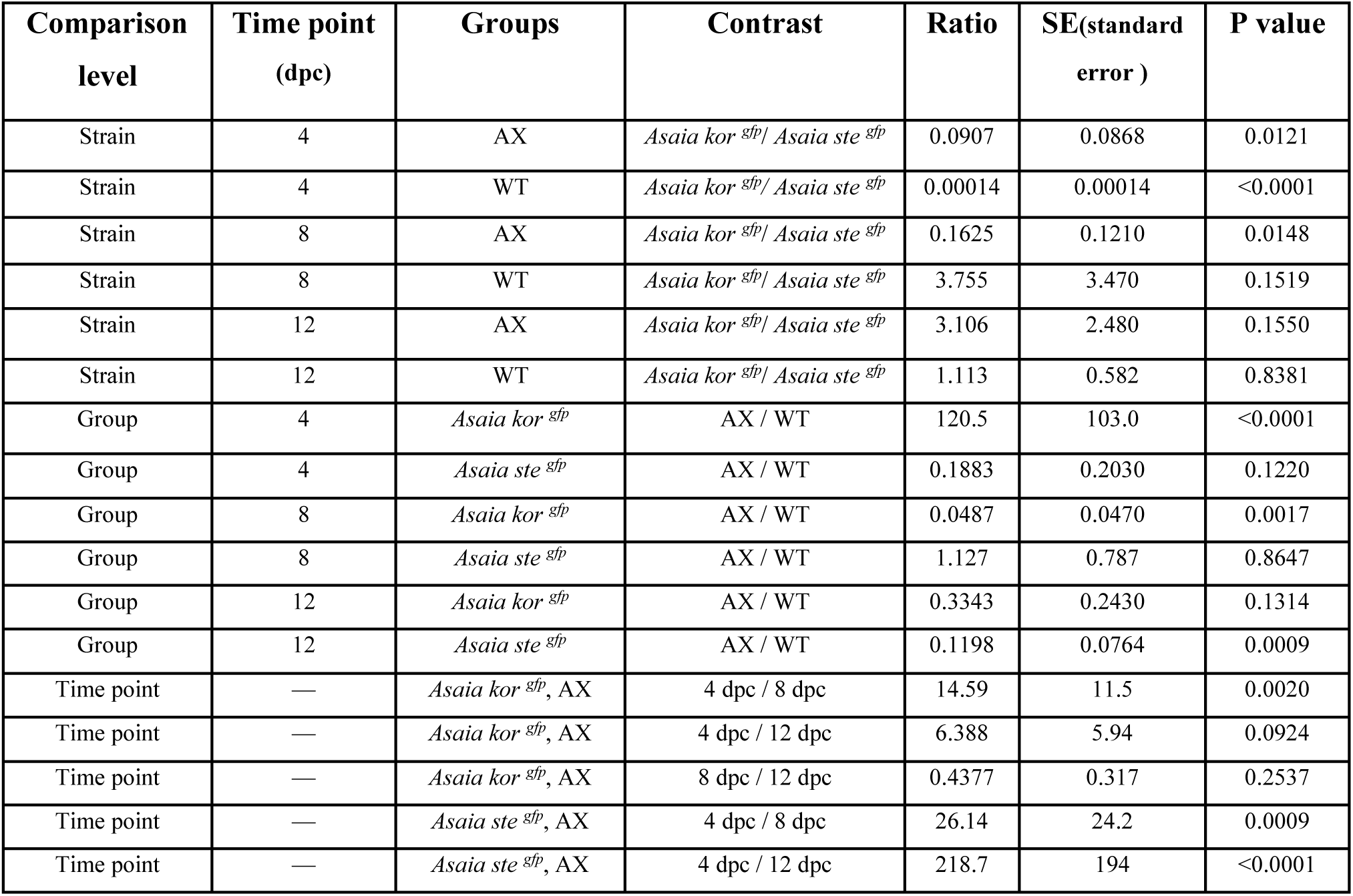
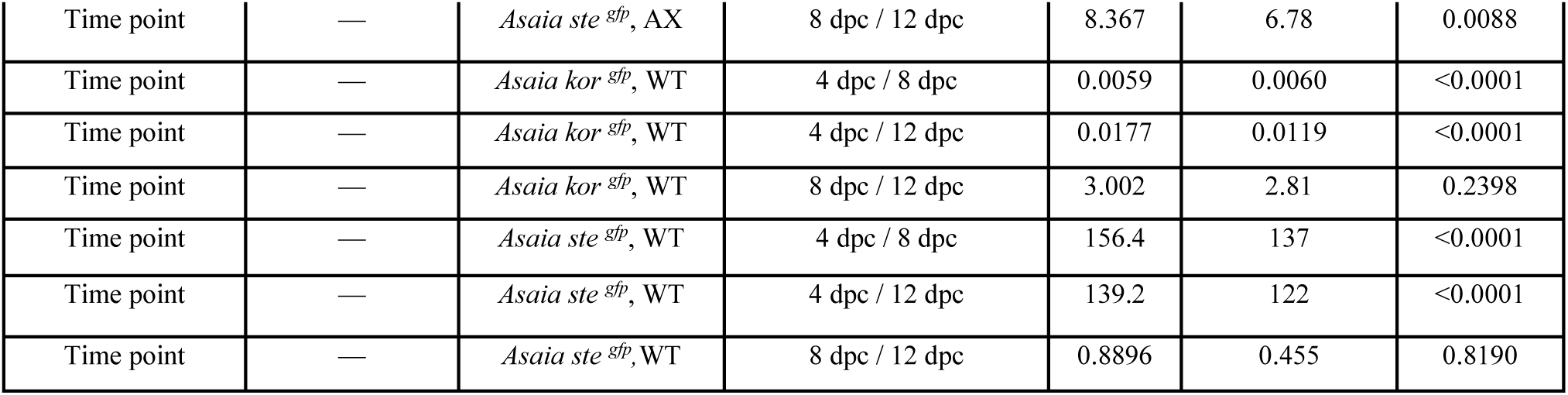
Pairwise comparisons of *Asaia* strain, axenic status, and time point on mosquito colonization in midgut.

| Comparison level | Time point (dpc) | Groups | Contrast | Ratio | SE(standard error ) | P value |
| --- | --- | --- | --- | --- | --- | --- |
| Strain | 4 | AX | *Asaia kor*<sup>gfp</sup> / *Asaia ste*<sup>gfp</sup> | 0.0907 | 0.0868 | 0.0121 |
| Strain | 4 | WT | *Asaia kor*<sup>gfp</sup> / *Asaia ste*<sup>gfp</sup> | 0.00014 | 0.00014 | <0.0001 |
| Strain | 8 | AX | *Asaia kor*<sup>gfp</sup> / *Asaia ste*<sup>gfp</sup> | 0.1625 | 0.1210 | 0.0148 |
| Strain | 8 | WT | *Asaia kor*<sup>gfp</sup> / *Asaia ste*<sup>gfp</sup> | 3.755 | 3.470 | 0.1519 |
| Strain | 12 | AX | *Asaia kor*<sup>gfp</sup> / *Asaia ste*<sup>gfp</sup> | 3.106 | 2.480 | 0.1550 |
| Strain | 12 | WT | *Asaia kor*<sup>gfp</sup> / *Asaia ste*<sup>gfp</sup> | 1.113 | 0.582 | 0.8381 |
| Group | 4 | *Asaia kor*<sup>gfp</sup> | AX / WT | 120.5 | 103.0 | <0.0001 |
| Group | 4 | *Asaia ste*<sup>gfp</sup> | AX / WT | 0.1883 | 0.2030 | 0.1220 |
| Group | 8 | *Asaia kor*<sup>gfp</sup> | AX / WT | 0.0487 | 0.0470 | 0.0017 |
| Group | 8 | *Asaia ste*<sup>gfp</sup> | AX / WT | 1.127 | 0.787 | 0.8647 |
| Group | 12 | *Asaia kor*<sup>gfp</sup> | AX / WT | 0.3343 | 0.2430 | 0.1314 |
| Group | 12 | *Asaia ste*<sup>gfp</sup> | AX / WT | 0.1198 | 0.0764 | 0.0009 |
| Time point | — | *Asaia kor*<sup>gfp</sup>, AX | 4 dpc / 8 dpc | 14.59 | 11.5 | 0.0020 |
| Time point | — | *Asaia kor*<sup>gfp</sup>, AX | 4 dpc / 12 dpc | 6.388 | 5.94 | 0.0924 |
| Time point | — | *Asaia kor*<sup>gfp</sup>, AX | 8 dpc / 12 dpc | 0.4377 | 0.317 | 0.2537 |
| Time point | — | *Asaia ste*<sup>gfp</sup>, AX | 4 dpc / 8 dpc | 26.14 | 24.2 | 0.0009 |
| Time point | — | *Asaia ste*<sup>gfp</sup>, AX | 4 dpc / 12 dpc | 218.7 | 194 | <0.0001 |
| Time point | — | *Asaia ste*<sup>gfp</sup>, AX | 8 dpc / 12 dpc | 8.367 | 6.78 | 0.0088 |
| Time point | — | *Asaia kor*<sup>gfp</sup>, WT | 4 dpc / 8 dpc | 0.0059 | 0.0060 | <0.0001 |
| Time point | — | *Asaia kor*<sup>gfp</sup>, WT | 4 dpc / 12 dpc | 0.0177 | 0.0119 | <0.0001 |
| Time point | — | *Asaia kor*<sup>gfp</sup>, WT | 8 dpc / 12 dpc | 3.002 | 2.81 | 0.2398 |
| Time point | — | *Asaia ste*<sup>gfp</sup>, WT | 4 dpc / 8 dpc | 156.4 | 137 | <0.0001 |
| Time point | — | *Asaia ste*<sup>gfp</sup>, WT | 4 dpc / 12 dpc | 139.2 | 122 | <0.0001 |
| Time point | — | *Asaia ste*<sup>gfp</sup>, WT | 8 dpc / 12 dpc | 0.8896 | 0.455 | 0.8190 |

**Table S3.** Pairwise comparisons of *Asaia* strain, axenic status, and time point on mosquito colonization in carcasses.

| Comparison level | Time point (dpc) | Groups | Contrast | Ratio | SE(standard error ) | P values |
| --- | --- | --- | --- | --- | --- | --- |
| Strain | 4 | AX | *Asaia kor*<sup>gfp</sup> / *Asaia ste*<sup>gfp</sup> | 1.143 | 0.905 | 0.8662 |
| Strain | 4 | WT | *Asaia kor*<sup>gfp</sup> / *Asaia ste*<sup>gfp</sup> | 1.126 | 1.190 | 0.9107 |
| Strain | 8 | AX | *Asaia kor*<sup>gfp</sup> / *Asaia ste*<sup>gfp</sup> | 0.2073 | 0.1520 | 0.0317 |
| Strain | 8 | WT | *Asaia kor*<sup>gfp</sup> / *Asaia ste*<sup>gfp</sup> | 0.0759 | 0.0648 | 0.0025 |
| Strain | 12 | AX | *Asaia kor*<sup>gfp</sup> / *Asaia ste*<sup>gfp</sup> | 0.8474 | 0.6380 | 0.8258 |
| Strain | 12 | WT | *Asaia kor*<sup>gfp</sup> / *Asaia ste*<sup>gfp</sup> | 1.513 | 1.140 | 0.5818 |
| Group | 4 | *Asaia kor*<sup>gfp</sup> | AX / WT | 0.2810 | 0.2900 | 0.2186 |
| Group | 4 | *Asaia ste*<sup>gfp</sup> | AX / WT | 0.2768 | 0.2120 | 0.0943 |
| Group | 8 | *Asaia kor*<sup>gfp</sup> | AX / WT | 9.040 | 7.80 | 0.0108 |
| Group | 8 | *Asaia ste*<sup>gfp</sup> | AX / WT | 3.308 | 2.54 | 0.1194 |
| Group | 12 | *Asaia kor*<sup>gfp</sup> | AX / WT | 0.0574 | 0.0430 | 0.0001 |
| Group | 12 | *Asaia ste*<sup>gfp</sup> | AX / WT | 0.1024 | 0.0772 | 0.0025 |
| Time point | — | *Asaia kor*<sup>gfp</sup>, AX | 4 dpc / 8 dpc | 5.561 | 4.47 | 0.0328 |
| Time point | — | *Asaia kor*<sup>gfp</sup>, AX | 4 dpc / 12 dpc | 44.42 | 32.6 | <0.0001 |
| Time point | — | *Asaia kor*<sup>gfp</sup>, AX | 8 dpc / 12 dpc | 7.989 | 6.16 | 0.0141 |
| Time point | — | *Asaia ste*<sup>gfp</sup>, AX | 4 dpc / 8 dpc | 1.009 | 0.722 | 0.9902 |
| Time point | — | *Asaia ste*<sup>gfp</sup>, AX | 4 dpc / 12 dpc | 32.94 | 24.6 | <0.0001 |
| Time point | — | *Asaia ste*<sup>gfp</sup>, AX | 8 dpc / 12 dpc | 32.65 | 24.2 | <0.0001 |
| Time point | — | *Asaia kor*<sup>gfp</sup>, WT | 4 dpc / 8 dpc | 178.9 | 205 | <0.0001 |
| Time point | — | *Asaia kor*<sup>gfp</sup>, WT | 4 dpc / 12 dpc | 9.069 | 9.68 | 0.0388 |
| Time point | — | *Asaia kor*<sup>gfp</sup>, WT | 8 dpc / 12 dpc | 0.0507 | 0.0436 | 0.0011 |
| Time point | — | *Asaia ste*<sup>gfp</sup>, WT | 4 dpc / 8 dpc | 12.06 | 8.89 | 0.0022 |
| Time point | — | *Asaia ste*<sup>gfp</sup>, WT | 4 dpc / 12 dpc | 12.19 | 9.09 | 0.0022 |
| Time point | — | *Asaia ste*<sup>gfp</sup>, WT | 8 dpc / 12 dpc | 1.011 | 0.724 | 0.9883 |

**Table S4.**
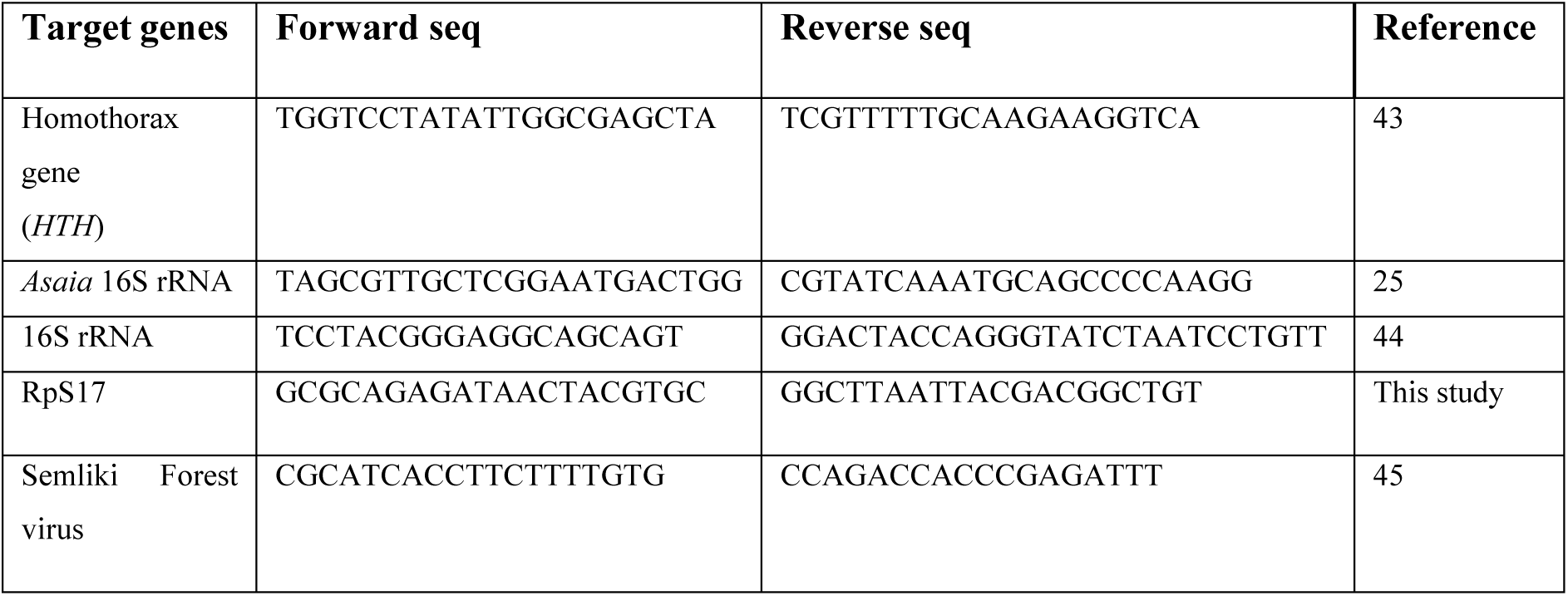
List of primers used in this study.

